# Tracking community change via network coherence

**DOI:** 10.1101/2025.11.04.686602

**Authors:** Alexandre Fuster-Calvo, Gracielle T. Higino, Christine Parent, Dominique Caron, Francis Banville, François Massol, F. Guillaume Blanchet, Katherine Hébert, Laura Pollock, Luigi Maiorano, Paulo R. Guimarães, Pablo Silva, Wilfried Thuiller, Dominique Gravel

## Abstract

Understanding how ecological communities respond to environmental change remains a key challenge for biodiversity monitoring. To characterize such responses, we need tools that capture how coherently species respond across a community, and to predict their consequences, we must account for ecological interactions. We first introduce the Ecological Coherence (EC) framework, which describes how species’ co-responses are structured within a community. Building on this foundation, we extend it to Ecological Network Coherence (ENC), which embeds co-responses within the network of interactions by restricting them to interacting species. Both are expressed through two complementary representations: a response correlation matrix and the distribution of its values. The first can reveal aspects such as coherent or incoherent modules and the roles species play in shaping coherence, whereas the second provides a profile whose shape may serve as an early-warning indicator of instability. These can be applied to both intrinsic responses (environmental performance) and realized responses (abundance dynamics), derived from currently available monitoring data. We illustrate this approach in two empirical systems: a tropical pollination network, where interacting mutualists were more coherent in their temperature responses than the broader community, and a marine food web, where coherence in abundance trends shifted during collapse. Using a Lotka–Volterra model, we further show that ENC distributions with higher variance—reflecting stronger positive and negative co-responses—increase the risk of instability or amplification in dynamics. We also find that species influential in both the correlation matrix and the interaction matrix are key drivers of major dynamic shifts. These results point to the importance of further exploring ENC distributions as potential early-warning indicators of ecological disruption.

## A need for characterizing network-level responses to environmental change

A central challenge in biodiversity monitoring is to capture collective signals of communities that may precede drastic shifts in structure, composition, and function (Berg et al. 2010, Magurran et al. 2010, Van der Putten et al. 2010, Trisos et al. 2020). One key aspect of such signals is species co-responses—the extent to which species track environmental change similarly or divergently. The insurance hypothesis provides a strong theoretical foundation that links this collective signal to stability: differential environmental responses among species, rooted in ecological trade-offs, generate asynchrony that buffers or even enhances ecosystem functioning (Yachi & Loreau 1999, Loreau & de Mazancourt 2008). This logic underpins two influential empirical frameworks: *Response diversity*, which emphasizes differences in species’ performance-environment relationships within functional groups, which allow some species to compensate for others under change (Elmqvist et al. 2003, Mori et al. 2013, Ross et al. 2023); and *Community synchrony*, which quantifies correlations in population fluctuations, showing that asynchrony stabilizes aggregate biomass while synchrony amplifies variability (Loreau & de Mazancourt 2008, Thibaut & Connolly 2013).

A key limitation of these frameworks, however, is that they largely ignore interaction structure (Loreau et al. 2021, Windsor et al. 2023). In real ecosystems, networks of interactions filter how co-responses occur, amplifying or buffering them and transmitting their effects across the system (Bellard et al. 2012, Harvey et al. 2017, Paine et al. 2018, Guimarães 2020; Cosmo et al. 2023, Zelnik et al. 2024). For example, an incoherent response in temperature-linked phenology can cause fish to spawn before phytoplankton blooms, producing a mismatch in larval feeding and resource availability (match–mismatch hypothesis; Cushing 1990, Asch et al. 2019). In mutualistic networks, coherent responses can preserve interaction persistence, whereas incoherent responses can destabilize links (Bartomeus et al. 2011, Cruz et al. 2023).

This gap is increasingly recognized. Emerging studies show that the stabilizing or destabilizing consequences of coherence depend strongly on network architecture and species’ roles within it (O’Connor et al. 2017, Viviani et al. 2019, Danet et al. 2021; Srednick & Swearer 2024). In other words, who responds together matters as much as how coherently they respond. To track and anticipate (sensu Maris et al. 2018) community dynamics under global change, we therefore need an integrated perspective that combines species’ co-responses to the environment with the architecture of their ecological networks (Windsor et al. 2023, Mendoza & Araújo 2025). Such a perspective would allow us to ask: does the way co-responses are arranged within the network structure affect community dynamics in predictable ways?

Here, we address this gap through Ecological Network Coherence (ENC), a framework that captures the variability of co-responses among interacting species. ENC builds on Ecological Coherence (EC), which first characterizes how co-responses are distributed within a community, by overlaying this information onto the network structure to capture variation among interacting species. Both the community-level (EC) and network-level (ENC) perspectives can be represented through two statistical objects: a correlation matrix and the distribution of its values, which can be derived from diverse response measures available in existing biomonitoring data. We illustrate this approach in two empirical systems: a tropical pollination network, where ENC revealed coherent thermal responses, and a collapsed marine food web, where ENC captured reorganization in coherence structure. We then use a Lotka–Volterra formulation and simulations to explore the potential consequences of different ENC distributions on community dynamics. Our results suggest that distributions with stronger coherence can increase instability, and that species with high coherence and high network centrality may act as key amplifiers or buffers of system-wide change. These findings motivate further exploration of ENC distributions as the basis for developing indicators to anticipate disruptive dynamics and abrupt ecological transitions from existing monitoring data.

## Ecological Network Coherence (ENC)

Conceptually, Ecological Coherence (EC) refers to how co-responses are structured within a community, whereas Ecological Network Coherence (ENC) extends this view to how co-responses are structured within a network of interactions. Here, coherence is used as a meta-concept that broadly refers to correlations in species’ responses—regardless of response type or the method used to quantify them. Formally, both EC and ENC are expressed through two complementary statistical objects: a correlation matrix across all species pairs (for EC) or restricted to interacting species (for ENC), and the distribution of the resulting values (Figure 1).

**Figure 1.**
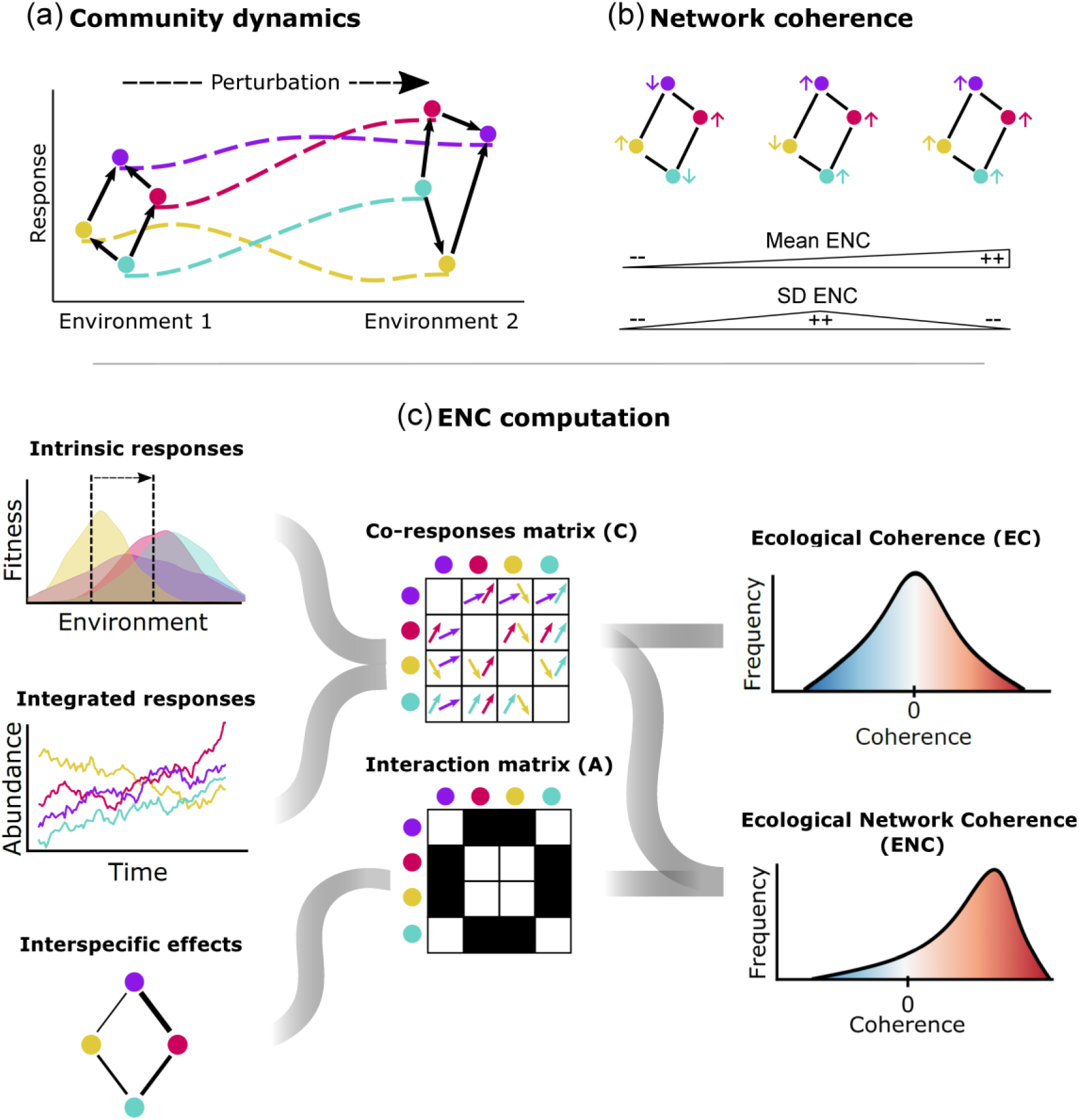
Conceptual framework of Ecological Coherence (EC) and Ecological Network Coherence (ENC). (a) Community dynamics emerge from the combined influence of species’ responses to the environment (dashed lines) and their ecological interactions (solid arrows). Interacting species may show varying degrees of coherence in their responses, either reinforcing or offsetting environmental effects. In turn, environmental coherence can influence the strength or direction of species interactions. These feedbacks shape community-level changes in abundance, distribution, and composition. (b) ENC quantifies how aligned or opposed the environmental responses are between interacting species. It captures both the sign (positive or negative coherence) and magnitude (weak to strong) of pairwise co-responses within the interaction network. In this framework, the interaction structure remains fixed. At maximum mean ENC, all interacting partners exhibit the same response to the environment, whereas at minimum mean ENC, complete opposite responses. (c) ENC can be estimated from widely available biodiversity monitoring data, including abundance time series (integrated responses) or species’ environmental sensitivities (intrinsic responses). Pairwise correlations or covariances in species’ responses form a co-response matrix (𝐶), whose distribution defines the Ecological Coherence (EC) profile for the community. Filtering this matrix through a binary interaction matrix (𝐴) yields the 𝐶_𝐴_ matrix (not shown), isolating co-responses among interacting species. The distribution of 𝐶_𝐴_values defines the ENC distribution, capturing the direction and strength of environmental co-responses within the network.

We define a community co-response matrix 𝐶, where each element 𝐶_𝑖𝑗_represents the pairwise correlation between the responses of species 𝑖 and 𝑗. Responses may be intrinsic, such as functional traits or performance measures that directly affect fitness — what response diversity theory terms response traits (Suding et al. 2008)—and that capture species’ fundamental environmental sensitivities. Alternatively, responses may be realized, such as abundance or biomass dynamics, which integrate these intrinsic responses, effects of species interactions, density dependence, and stochasticity. The matrix is symmetric by definition (𝐶_𝑖𝑗_ = 𝐶_𝑗𝑖_) and can be extended to a higher-dimensional structure when intrinsic responses to multiple variables are considered. For simplicity, we focus here on the single-variable case. The structure of the 𝐶 matrix contains ecologically meaningful information. It can identify clusters of species with more negatively or positively correlated responses (coherent or incoherent modules), and quantify how individual species contribute to community coherence, such as through the average or evenness of their correlations with others.

The distribution of correlation values in 𝐶, which we term *EC distribution*, provides a summary of how coherence is structured at the community level. Unlike previous approaches that collapse co-responses into a single index (Houlahan et al. 2007, Thibaut et al. 2012, Houlahan et al. 2018, Valencia et al. 2020, Ross et al. 2023, Tsang et al. 2023, Granzotti et al. 2024), the EC distribution characterizes their variability and shape: a normal distribution centered on zero indicates unstructured responses, skewed distributions reflect overall coherence or divergence, and U-shaped or heavy-tailed distributions suggest the presence of tightly coupled modules or a coexistence of highly coherent and highly divergent pairs.

However, like existing coherence frameworks, EC does not account for the interaction network. Yet predicting community-level responses requires considering how environmental effects propagate through species interactions. ENC addresses this gap by filtering EC through the network of interactions to retain only co-responses among interacting species. Specifically, we use the community’s adjacency matrix 𝐴, a symmetric matrix where 𝐴_𝑖𝑗_ = 1 if species 𝑖 and 𝑗 interact, and 𝐴_𝑖𝑗_ = 0 otherwise. By applying the Hadamard (element-wise) product between the community response-correlation matrix 𝐶 and the adjacency matrix 𝐴, we obtain the network response-correlation matrix 𝐶_𝐴_:

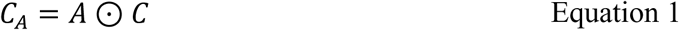

This filtered matrix retains only the co-responses among interacting species, thereby capturing the network-filtered structure of community coherence. The distribution of the non-zero elements in 𝐶_𝐴_ forms the *ENC distribution*. Together, the 𝐶_𝐴_matrix and the resulting ENC distribution provide tools to evaluate co-responses in the context of interactions, allowing us to ask: how coherent are networks to environmental change? they also enable more detailed analyses, such as quantifying coherence within modules of interacting species, assessing whether it is concentrated in particular network regions such as the core or periphery, and comparing how coherence varies among species with different network roles. More importantly, because networks modulate how response coherence translates into community dynamics (Haynes et al. 2009, Montoya et al. 2009, Guimarães et al. 2017, Eschenbrenner & Thébault 2023, Danet et al. 2024, Terry et al. 2025), ENC raises the question of whether ENC distributions could serve as early indicators of instability and vulnerability.

## Tracking ENC

EC and ENC are flexible to the choice of method used to quantify co-responses. For example, synchrony in population dynamics can be estimated through pairwise correlations, variance ratio metrics, or hierarchical modeling approaches (Ives 1995, Gouhier et al. 2010, Hébert 2024). Similarly, intrinsic sensitivities can be derived from functional traits, performance-environment curves, or species distribution models. Our intention is not to prescribe a single metric, but to provide a generalizable framework in which different approaches can be compared, and their complementarity explored. We propose here to focus on two response types that can be broadly estimated from biodiversity monitoring data: abundance trends modeled with generalized additive models (GAMs), and environmental sensitivities derived from species distribution models (SDMs). These responses are accessible for diverse taxa and ecosystems, enabling coherence to be applied across systems and scales (see section Tracking ENC).

Given the urgency of global change, it is essential to develop actionable tools that can be applied to data that biodiversity monitoring programs already generate. Here, we propose to compute ENC specifically from three types of data that are the most widely available across taxa and ecosystems: occurrence records (e.g., GBIF) and abundance time series (e.g., BIOTIME; Dornelas et al. 2018) to approximate species’ responses, and binary interactions (e.g., GLOBI; Poelen et al. 2014, Mangal; Poisot et al. 2016) to approximate ecological networks. The choice of response type depends on the ecological question, while binary networks provide the most consistent information across systems, with robust predictive frameworks now available to extend their coverage across space and time (Massol et al. 2011, Gravel et al. 2013, Poisot et al. 2015, Cazelles et al. 2016, Gravel et al. 2018, Albouy et al. 2019). These choices make ENC readily computable across systems. Below, we outline practical ways to derive coherence in responses from occurrence and abundance data, construct EC matrices and their distributions, and, when combined with interaction networks, estimate the ENC distribution.

### Coherence in sensitivity to the environment

Intrinsic environmental responses—those expressed independently of species interactions— are central to understanding community-level dynamics. The response diversity framework has been influential in this regard. It quantifies the degree of complementarity in species’ functional response traits, i.e., phenotypic characteristics that affect performance and fitness (Suding et al. 2008), by measuring their dispersion in multidimensional trait space (Cadotte et al. 2011). These metrics capture the idea that heterogeneous trait combinations underlie differential species responses to environmental change, and thus stabilize community functioning.

Yet this trait-based approach has important limitations. Many commonly measured traits (e.g. leaf or root characteristics; Sasaki et al. 2019) have uncertain or indirect links to performance under environmental gradients (Bartomeus et al. 2018), and several traits often need to be measured simultaneously to approximate fitness responses. Empirical data are also uneven, being mostly concentrated in plant communities (Ross et al. 2023) or specific animal groups like birds (Hordley et al. 2021), with far less coverage for multi-taxa, trophically diverse systems. Finally, most implementations summarize coherence variation into a single index such as functional dispersion (FDis; Laliberté et al. 2010) or similar measures (Correia et al. 2018), and mask the underlying structure of responses.

A more compelling strategy is to focus directly on environmental sensitivities, i.e, how species’ performance changes along environmental gradients. Ross et al. (2023) propose fitting species-specific performance–environment curves (e.g. biomass or growth vs. temperature) with flexible models, then using the distribution of their derivatives as a direct measure of response diversity. This approach, however, requires detailed data on demographic rates, growth, or carrying capacities—data still scarce beyond a few taxa. New initiatives such as GlobTherm (Bennett et al. 2018) are beginning to compile thermal tolerance curves for multiple taxa, but coverage still remains highly incomplete across ecosystems and trophic guilds.

For broader applications, especially when summarizing responses across diverse communities, species distribution models (SDMs) offer a practical alternative (Gómez-Ruiz & Lacher 2019). By fitting occurrence data to environmental predictors, SDMs estimate species’ realized niches and provide response curves to key abiotic gradients. Despite limitations such as sampling bias and environmental collinearity (Wiens et al. 2009), SDMs have already been used to project community disruption via range shifts and guild mismatches (Selden et al. 2018, Trisos et al. 2020). Within the ENC framework, they can be leveraged to quantify how species’ environmental responses align within interaction networks.

To operationalize this, species’ sensitivities can be derived as the derivative of their response function with respect to an environmental variable, evaluated at each grid cell in space (𝜕𝑝_𝑖_(𝐸)⁄𝜕𝐸). Correlating these spatially explicit sensitivities across species yields the co-response matrix 𝐶, whose distribution (EC) summarizes community coherence to the environment within a given spatial range. Filtering 𝐶 by the interaction network isolates co-responses among interacting species, producing the ENC distribution. This distribution reveals whether interacting species respond coherently or divergently to environmental gradients, and potentially providing early signals of mismatch, vulnerability, or stabilization potential.

In an Atlantic Rainforest pollination network (see Box 1), we modeled species’ responses to temperature as truncated normal distributions fitted to the environmental values at known occurrences. We derived species’ response curves to mean temperature from their occurrence data using fitted probability density functions. For each species, we then computed the derivative of its response curve with respect to temperature at each spatial cell across the study area. Correlating these derivative values across species yielded the 𝐶 matrix, which summarizes how similarly species respond to temperature across space. This analysis revealed that species pairs with documented interactions (ENC) tend to exhibit more consistently positive co-responses than the community at large (EC), suggesting that ecological coherence is stronger among interacting species. Moreover, some highly connected pollinators showed strong coherence, identifying them as potential stabilizers of network-wide coherence.

#### Box 1. Case study 1: ENC in an Atlantic Rainforest pollination network

We assessed the Ecological Network Coherence (ENC) of a pollination network from the Atlantic Rainforest of South-Eastern Brazil. The bipartite network comprised 23 bromeliad species and 22 pollinators, including hummingbirds, bees, and butterflies, sourced from the Interaction Web Database (Interaction Web DataBase, n.d.; Varassin & Sazima, 2012).

To quantify environmental responses, we assembled species occurrence records from the Global Biodiversity Information Facility (GBIF), filtering records to fall within the Atlantic Rainforest biome. Occurrence data were cleaned to remove duplicates, misidentified or ambiguous records, and species represented by a single locality. For each valid occurrence point, we extracted the mean annual temperature (WorldClim BIO1) at approximately 4.6 km resolution, which we used as the focal environmental variable (Figure 2a).

**Figure 2.**
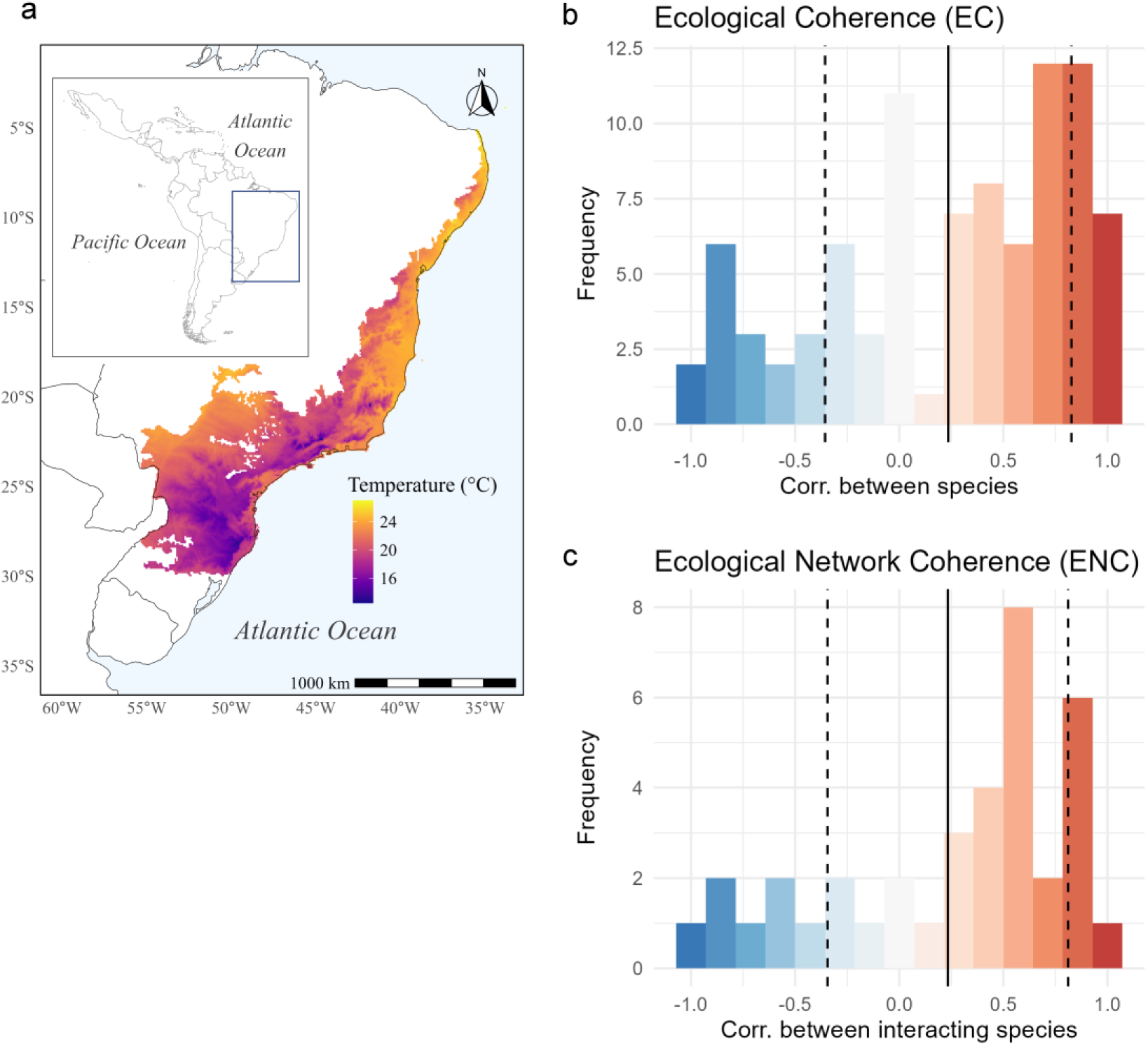
EC and ENC in an Atlantic Rainforest pollination network. (a) Mean annual temperature across the Brazilian Atlantic Rainforest ecoregion, used to compute species’ environmental responses. (b) Ecological Coherence (EC) represents the distribution of pairwise Spearman correlations between species’ environmental response slopes (i.e., first derivatives of species occurrence curves with respect to temperature), computed across shared spatial cells. (c) Ecological Network Coherence (ENC) filters these pairwise correlations to retain only those between species that are directly connected in the pollination network.

For each species, we approximated its environmental response using a truncated normal probability density function (PDF) fitted to its distribution of observed temperature values (defined by the species’ empirical mean, standard deviation, and range; Figure S1). We then computed the derivative of this PDF with respect to temperature at each occurrence point using numerical differentiation. These cell-level derivatives represent the instantaneous rate of change of occurrence likelihood with respect to temperature—a proxy for environmental response. To compute Ecological Coherence (EC), we calculated pairwise Spearman correlations between species’ derivative values across shared grid cells. This resulted in a complete species-by-species correlation matrix representing the similarity in environmental responses across the entire community.

To assess Ecological Network Coherence (ENC), we intersected the EC matrix with the pollination network, retaining only the correlation values for species pairs that were observed to interact. To reduce spurious values, we included only interactions for which both plant and pollinator had at least two shared spatial cells with derivative estimates, yielding a final dataset including 17 bromeliad and 9 pollinator species.

To assess whether observed EC and ENC distributions differed from chance, we randomized temperature values independently across species at each site, recalculating response curves and their derivatives, and repeating the process 100 times to generate null EC and ENC distributions. For each null, we computed the corresponding 𝐶 and 𝐶_𝐴_ matrices and compared the observed vs. null distributions using the Wasserstein distance, implemented via the *wasserstein1d* function from the *transport* R package (Schuhmacher et al., 2017). Significance was evaluated by comparing the mean observed-null distance to the distribution of null-null distances, yielding one-sided empirical p-values. Observed EC and ENC distributions differed significantly from null expectations (p < 0.05).

The distribution of Ecological Coherence (EC) across all species pairs was skewed toward positive values, indicating that many species tend to respond similarly to temperature gradients (Figure 2b). However, a substantial number of contrasting responses were also observed, potentially reflecting niche differentiation, spatial turnover, or asynchronous phenologies. A prominent secondary peak near zero correlation likely reflects species pairs with limited or no spatial overlap, where correlations are computed over very few shared cells, resulting in unstable or default values close to zero—an expected artifact in diverse communities with uneven spatial distributions. In contrast, the ENC distribution showed a skew toward positive values but the frequency of neutral or negative correlations was low (Figure 2c).

This suggests that ecological coherence is more consistently positive among interacting species compared to the community at large. This observation may arise through two complementary processes. First, environmental similarity may act as a filter, favoring interactions among species that respond similarly to climate. For example, through phenological synchrony or shared habitat preferences. Second, mutualistic interactions may reinforce coherence over time through coevolution, promoting the alignment of species’ environmental responses.

We also calculated degree centrality for each species from the unweighted bipartite network and quantified each species’ contribution to EC as the median of its pairwise correlation values across all other species in the community. The analyses revealed that most species showed positive median correlations in their environmental responses, indicating coordinated tracking of environmental change (Figure 3). Notably, high-centrality hummingbirds such as *Phaethornis spp*. and *Thalurania glaucopis* also exhibited strong coherence, suggesting that these species may act as stabilizing agents within the community by aligning interaction dynamics with shared environmental responses. In contrast, plant species generally exhibited lower degree centrality and more variable coherence. Notably, *Aechmea victoriana* showed strongly negative coherence, suggesting a heightened sensitivity to microhabitat conditions or a distinct ecological strategy.

**Figure 3.**
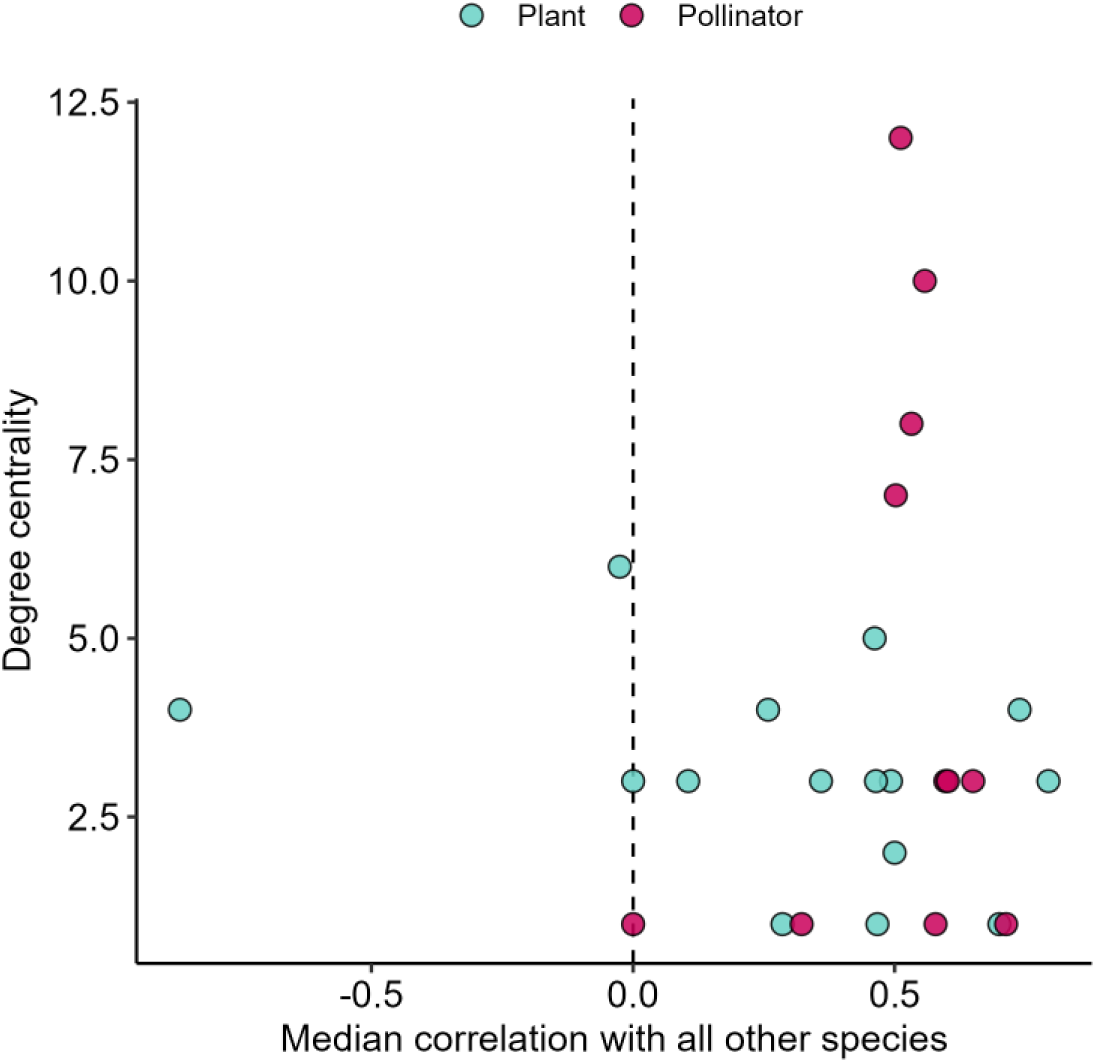
Species roles in ecological coherence to temperature and network structure. Relationships between species’ roles in the network (centrality) and their contribution to community coherence (i.e., median pairwise correlation of a species’ environmental response with that of all other species in the community).

### Coherence in fluctuations of abundances

Abundance dynamics integrate the combined effects of intrinsic environmental responses, biotic interactions, and stochasticity. This makes abundance time series a powerful but complex signal of community change. The synchrony-stability framework predicts that negative coherence—or anti-correlated dynamics—among species’ responses in abundance create compensatory dynamics that can stabilize aggregate community properties such as biomass or productivity (Loreau & de Mazancourt 2008). While this has strong support in horizontally structured or competitive communities, recent studies show it does not hold in more complex systems with vertical structure; instead, how synchrony distributes across functional groups, trophic levels, or influential species may play a greater role in shaping community dynamics (Viviani et al. 2019, Huang et al. 2020, Valencia et al. 2020, Danet et al. 2021, Rao et al. 2024, Srednick & Swearer, 2024). ENC may offer a synthetic representation of that complexity.

Within time series data also allows ENC to be tracked dynamically, identifying transitions such as the buildup or erosion of ENC distribution shapes before certain abrupt dynamics such as collapse emerge. They further link species’ trajectories to community dynamics by combining information on network position and coherence with other species, revealing potential amplifiers or buffers of change. Moreover, coherence can be examined at different temporal scales. For example, short-term year-to-year or seasonal fluctuations versus longer-term multi-decadal trends. These different frequency components may carry distinct ecological meanings and signal different types of risk: short-term synchrony may reflect common responses to immediate drivers such as climate anomalies (e.g. heat waves; Arimitsu et al. 2021), whereas coherence in long-term trends may foreshadow lasting reorganization of community structure.

Methodologically, abundance-based coherence can be estimated from time series using approaches ranging from year-to-year pairwise Pearson correlations in abundance change for short-term coherence (Lertzman-Lepofsky et al. 2025) to hierarchical dynamic GAMs with latent temporal predictors to recover long-term coherence (Gouhier et al. 2014). In our fish community case study (Box 2), we applied this later approach to infer biomass trajectories and compute 𝐶 matrices throughout its collapse. Filtering 𝐶 through the interaction network yielded the 𝐶_𝐴_matrix and ENC distribution over time, revealing a transition from structured coherence (pre-collapse), to fragmentation (collapse), to a simplified, tightly synchronized core (post-collapse), and identified taxa likely to drive or buffer community-level dynamics.

#### Box 2. Case study 2: ENC in a destabilized fish community

The Newfoundland shelf groundfish community (Figure S2) underwent significant restructuring during the 1980s-1990s following the near-collapse of key species like the Atlantic cod (*Gadus morhua*) due to intense fishing pressure and environmental changes (Pedersen et al. 2017). This collapse triggered cascading shifts in community composition and destabilized species dynamics.

To quantify changes in Ecological Coherence (EC) and Ecological Network Coherence (ENC), we analyzed annual biomass time series for 29 groundfish species across three periods: pre-collapse (1980-1989), collapse (1990-1999), and post-collapse (2000-2013). A Bayesian hierarchical model (Dynamic GAM; Clark & Wells 2023) was used to estimate species-specific biomass trends (see Appendix 1 for more details) and compute co-response matrices 𝐶, which capture pairwise correlations in annual fluctuations. For ENC, these matrices were filtered through a binary trophic interaction network 𝐴, sourced from GloBI (Poelen et al. 2014), to retain only co-responses between interacting species. This produced the 𝐶_𝐴_matrix, which represents the structure of coherence within the ecological network. Species’ contributions to coherence were summarized as their median correlation with other taxa, and network influence was measured by degree centrality.

To assess whether observed EC and ENC distributions deviated from expectations under independent dynamics, we generated 100 null models per period by permuting each species’ time series independently and refitting the Bayesian model. As in Box 1, for each null, we computed the corresponding 𝐶 and 𝐶_𝐴_ matrices and compared the observed vs. null distributions using the Wasserstein distance. Significance was evaluated by comparing the mean observed-null distance to the distribution of null-null distances, yielding one-sided empirical p-values. In all periods, observed EC and ENC distributions differed significantly from null expectations (p < 0.05).

The EC distributions revealed a gradual structural shift in community-wide co-responses (Figure 4a). Pre-collapse, the distribution was U-shaped, with a predominance of strong positive and negative correlations, indicating polarized dynamics. During the collapse, moderate correlations increased and extreme ones became less dominant. Post-collapse, EC approximated a normal distribution centered on zero—still retaining some tails—reflecting a general weakening of co-responses.

**Figure 4.**
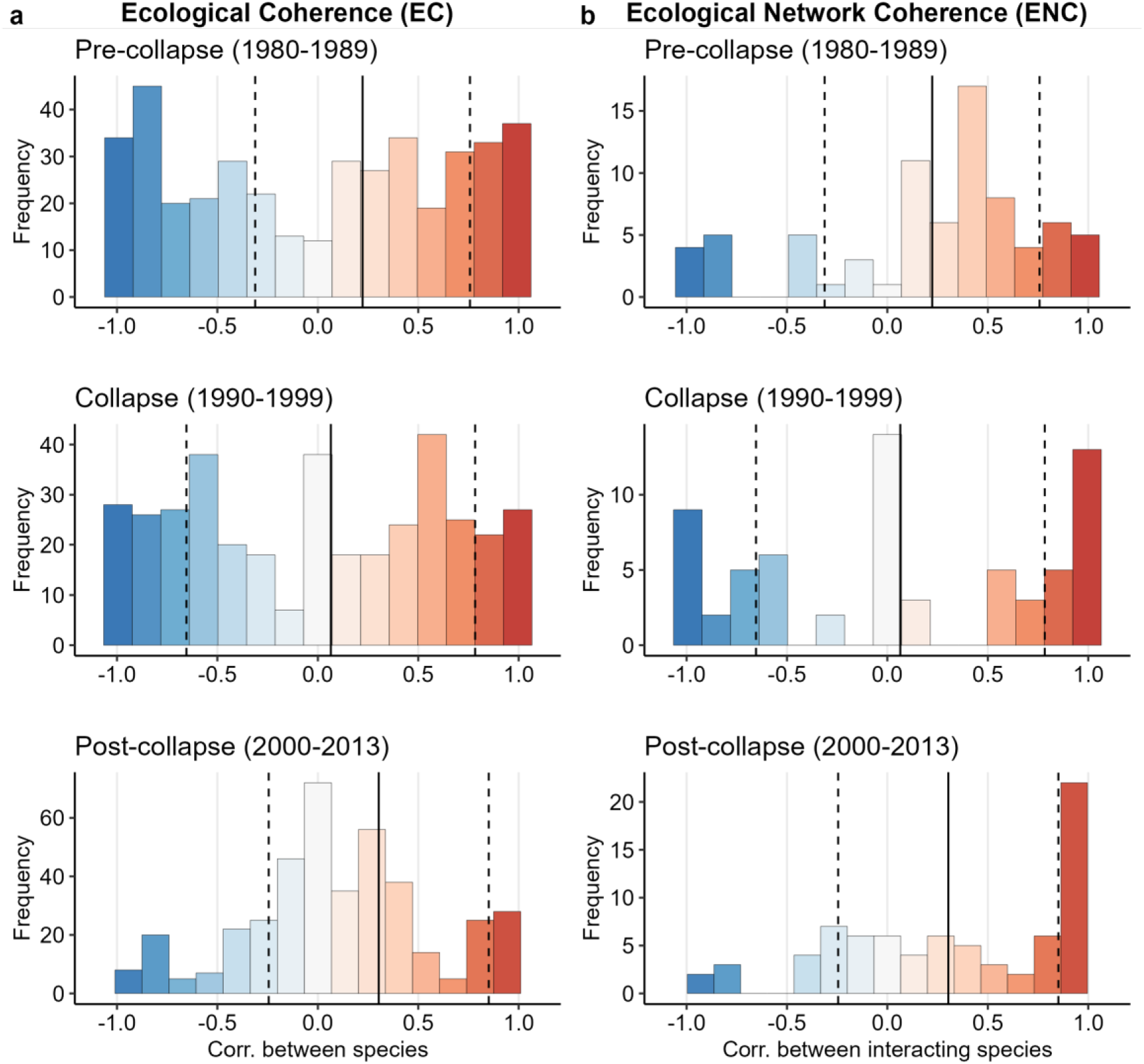
Ecological Coherence (EC) and Ecological Network Coherence (ENC) during the collapse and initial recovery of a groundfish food web in the Northwest Atlantic Shelf. (a) EC distributions showing the pairwise correlations in annual biomass fluctuations across all species in the community, and (b) the ENC distribution, restricting this view to interacting pairs. Solid vertical lines indicate the mean correlation; dashed lines represent one standard deviation from the mean. Distributions are shown for three periods: pre-collapse (1980–1989), collapse (1990–1999), and post-collapse (2000–2013).

ENC distributions refined this picture by revealing how coherence was structured within the interaction network (Figure 4b). Before the collapse, the ENC distribution was skewed toward moderate and strong positive co-responses, suggesting that coherence was concentrated within specific trophic modules. This skew also indicated a potential vulnerability, as such clustering could lead to over-synchronization and increase the risk of system-wide instability. During the collapse, the distribution became trimodal and more dispersed: many interacting species became weakly coupled, while others exhibited extreme synchrony or divergence, signaling a breakdown in the organization of interaction-mediated responses. In the post-collapse phase, a small group of tightly synchronized, interacting species dominated the network. This distribution suggests the emergence of a simplified core of functionally dominant species, likely reflecting trophic simplification, surrounded by more decoupled or functionally redundant taxa.

By comparing species’ contributions to coherence and their structural position in the network, we found that highly connected predators like cod were central contributors to ENC (Figure 5). Their sharp decline would not only have reduced top-down control but also disrupted the coherence of the network itself.

**Figure 5.**
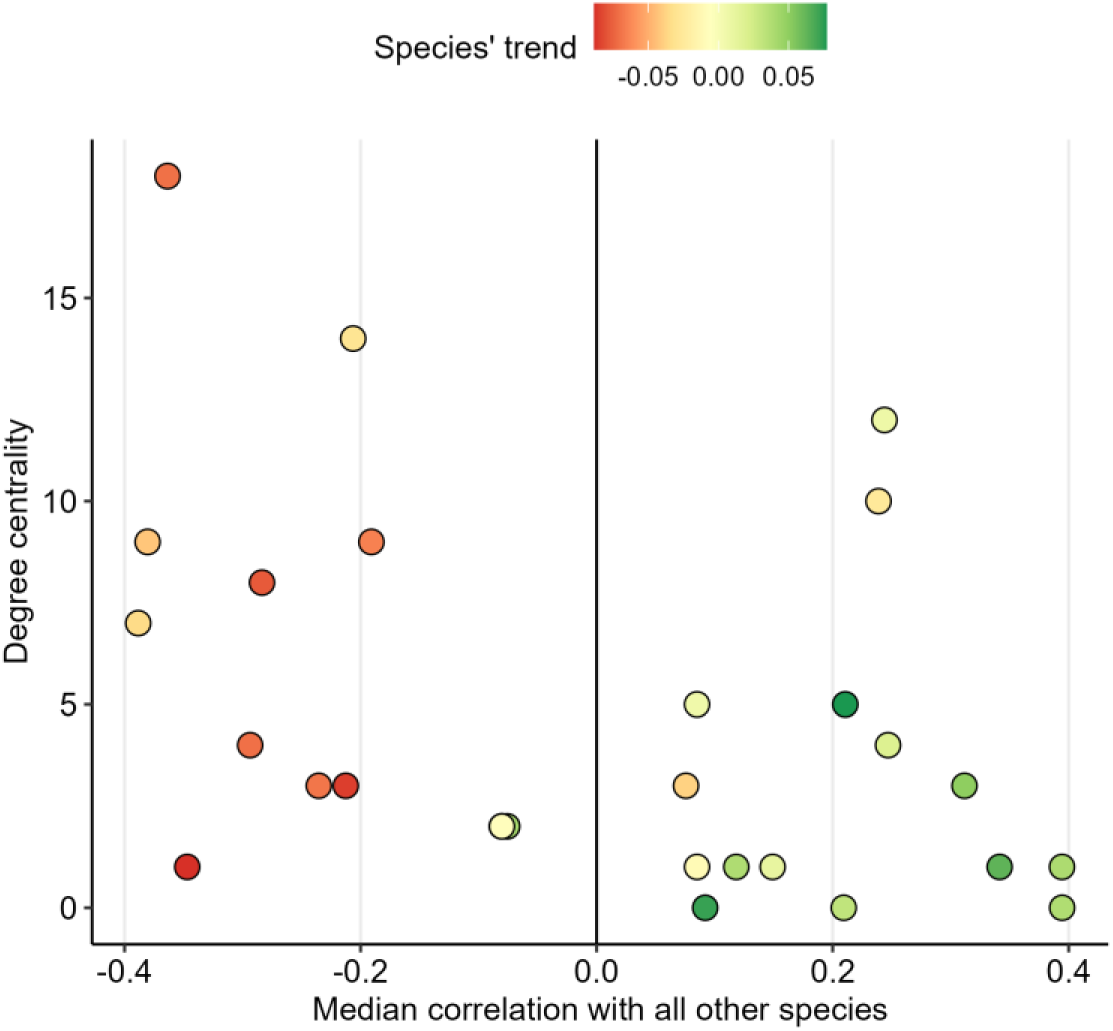
Species roles in ecological coherence and network structure during community collapse. Relationships between species’ roles in the network (centrality), their contribution to community coherence (i.e. median pairwise correlation of a species’ response with that of all other species in the community), and their average annual rate of change in biomass over the period from 1980 to 2013.

Together, EC and ENC provide complementary insights: EC revealed a general shift from structured extremes to weaker, more homogeneous synchrony. ENC, by contrast, highlighted how coherence is lost, reorganized, and re-concentrated among interacting species. This combination allows us to distinguish between diffuse shifts in synchrony and structural reorganization within the network.

#### Box 3. Simulating ENC effects on community stability

We first provided theoretical intuition for how co-responses can combine with a network of interactions to generate unstable community dynamics. Here, we extend this by presenting simulations that use a fixed interaction matrix 𝐴 together with different EC distributions, in order to explore how variation in co-response structure shapes system-level outcomes.

While natural ecosystems may exhibit diverse covariance distributions, our simulations were constrained to normal distributions centered on 0 due to the requirement of positive definiteness for generating valid correlation matrices. We used a generalized Lotka-Volterra model with eight species interacting in a fixed food web, represented by a quantitative interaction matrix 𝐴. This matrix was generated using a predefined connectance and interaction strength distribution (Figure 6a), and remained constant within each set of scenarios.

**Figure 6.**
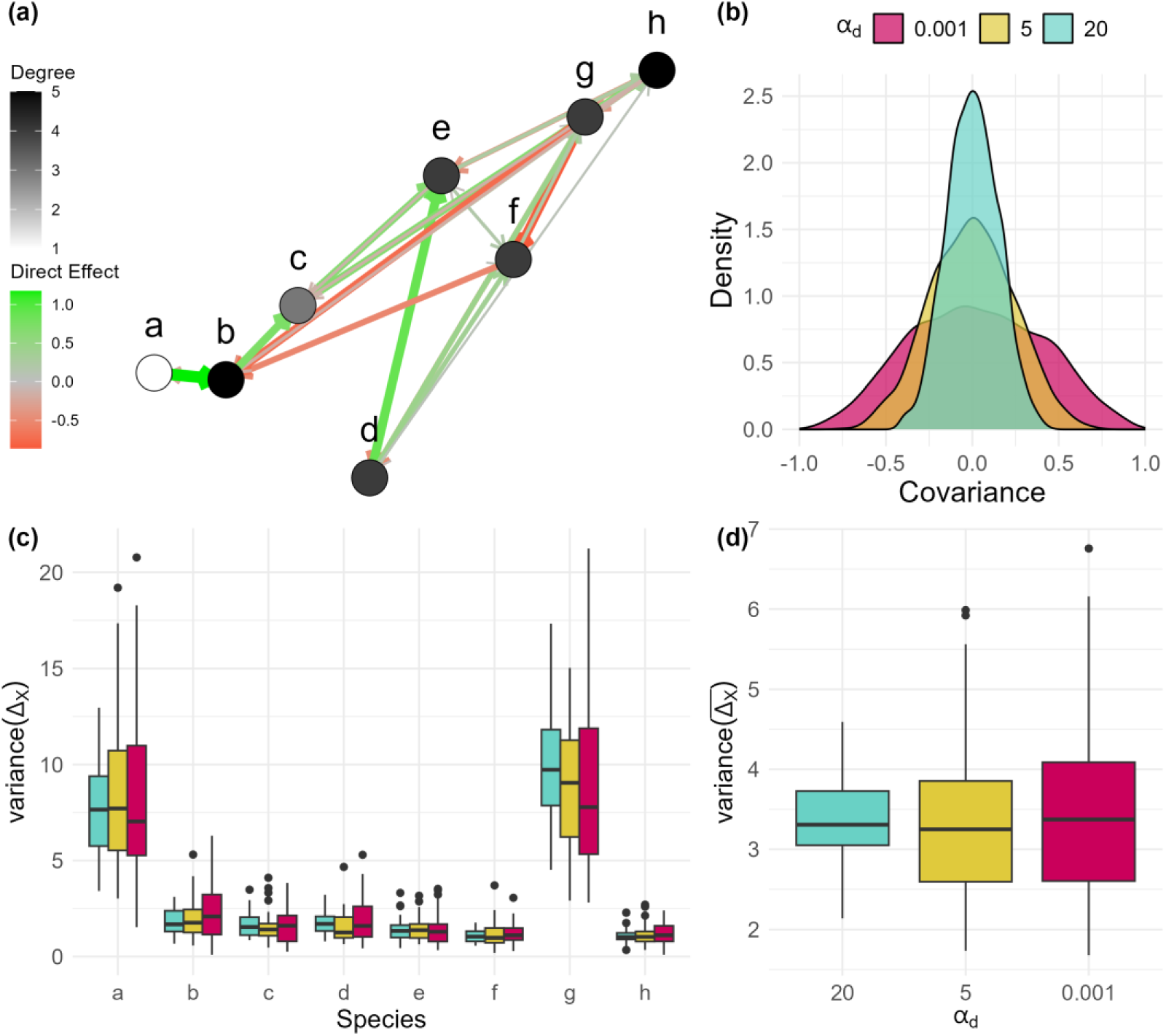
Consequences of EC distributions on community stability. (a) The fixed interaction matrix 𝐴 used across all scenarios, representing the food web structure. (b) Distributions of coherence values, i.e., all species co-responses, from all the co-response matrices 𝐶 generated for each investigated covariance structure. (c) Species-level variance in mean abundance change across simulations for different coherence structures. (d) Community-level variance in mean abundance change across simulations, illustrating the cumulative effects of varying coherence structures on overall community dynamics.

We also generated 50 different distributions of covariance of species responses using the *rcorrmatrix* function from the *clusterGeneration* R package (Qiu et al. 2015). This function samples a random correlation matrix 𝐶 controlled by a shape parameter 𝛼_𝑑_, controlling for the distribution of partial correlations: higher 𝛼_𝑑_ values yield weaker and more independent responses, while lower values produce stronger and more coherent co-responses (Figure 6b).

In each scenario, we ran 20 stochastic simulations of abundance dynamics across three phases: First, species intrinsic growth rates 𝑟 were set to counterbalance interactions (i.e., 𝑟 = 𝐴𝑋) at a random initial biomass, and the system was run to equilibrium; second, we sampled a perturbation Δ𝑟 ∼𝑁(0, ∑) where ∑ = 𝐶𝜎^2^, representing a change to species’ intrinsic growth rates. This structured perturbation encoded the hypothesized structure of co-responses; and third, the perturbed 𝑟 values were applied, and the system was again run to equilibrium. We recorded the change in species abundances (Δ𝑋) between pre- and post- perturbation equilibria.

We found that coherence structures with higher variance (i.e., with stronger co-responses corresponding to low 𝛼_𝑑_) increased the likelihood of cascading effects, leading to heightened variability in community dynamics (Figure 6d). The most specialized species (a), as well as a generalist involved in strong interactions (g), exhibited greater variance in abundance changes (Figure 6c).

### Bias and null models

Abundance and occurrence measures are subject to statistical bias susceptible to influence the distribution of co-responses. For instance, correlations between species’ log-abundances may be inflated by shared responses to unmeasured environmental drivers, or skewed by differences in time series length, variance, or prevalence (Shoemaker et al. 2022). Similarly, slopes derived from species distribution models (e.g., the derivative of the occurrence-environment curve) may be influenced by sampling intensity, environmental collinearity, or uneven geographic coverage (Soley-Guardia et al. 2024). Consequently, to interpret observed EC and ENC, they must be compared to null expectations. In our case studies, we generated null distributions by randomizing time series (for occurrence data; Box 1) or temperature values (for abundance data; Box 2) independently across species, allowing us to test whether observed coherence exceeded expectations under independent dynamics.

### Consequences on community stability

Coherence in a network may influence how a community responds to environmental change. For instance, a food web might remain functionally intact if most species shift similarly under warming, but lose trophic regulation if consumers and prey diverge in their responses. By capturing the structure of coherent and incoherent responses across a network, ENC may therefore serve as an indicator of potential instability and reorganization.

In this section, we derive baseline expectations to motivate future work on the consequences of ENC for community dynamics. To this end, we use an idealized Lotka–Volterra system in which coherence is defined over species’ intrinsic growth rates (𝑟)—a quantity only approximated by abundance time series or sensitivities to the environment derived from occurrence data (Webber et al. 2017, Osorio-Olvera et al. 2019). This yields an EC distribution, which we then combine with the community interaction matrix 𝐴, where its inverse 𝐵 = 𝐴^−1^summarizes the net direct and indirect effects of species on one another. Unlike ENC, we do not apply an explicit filtering step to restrict co-responses to directly interacting species. Instead, the framework assumes a fully connected system in which environmental variance propagates along both direct and indirect pathways (Martins et al. 2024). This should be seen not as ENC per se, but as a conceptual demonstration of how distributions of co-responses interact with network structure to shape community instability, and why ENC distribution profiles (e.g. clustered, skewed, polarized) may provide useful proxies for community vulnerability.

### Model and definitions

We are interested in documenting how co-responses and species interactions jointly affect a vector of community composition 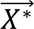. We assume a closed system (no immigration) undergoing Lotka-Volterra dynamics, with a fixed interaction network in which the set of interacting species remains constant. The abundance 𝑋_𝑖_ of species 𝑖 changes according to:

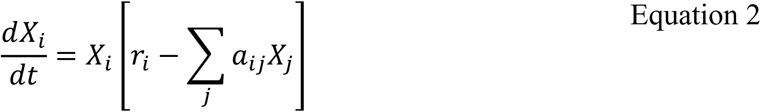

where 𝑟_𝑖_ is the intrinsic growth rate and 𝑎_𝑖𝑗_ is the interaction coefficient. At equilibrium, assuming all species persist (Novak et al. 2016):

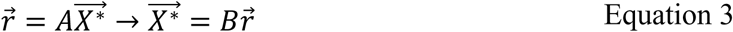

where 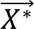 is the vector of equilibrium abundances, 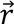 is the vector of intrinsic growth rates, 𝐴 is the interaction matrix, and 𝐵 = 𝐴^−1^. If a press perturbation 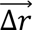 is applied to the system over a sufficiently long time, it shifts the equilibrium by:

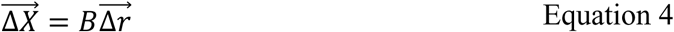

We are interested in documenting how a change in the environment affecting the intrinsic growth rate Δ𝑟 scales up and drives changes in abundance Δ𝑋. More specifically, we are seeking to determine if there is a relationship between matrix 𝐴, ecological coherence (here cov(Δ𝑟, Δ𝑟)), and the properties of the distribution of Δ𝑋. The expected change in abundance across the community 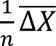 is:

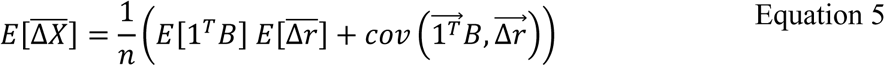

The first term reflects how the average environmental effect aligns with the network’s structure; the second term captures whether species with strong influence (high column sums in B) are also those most affected, amplifying system-wide change if these two align.

The variance in species 𝑖’s abundance change is:

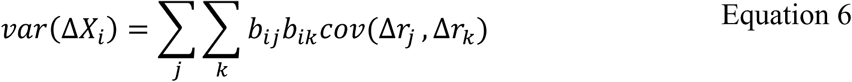

where 𝑏_𝑖𝑗_and 𝑏_𝑗𝑖_are elements of the matrix 𝐵 = 𝐴^−1^. Variance depends on two factors: (1) how strongly species 𝑖 is influenced by others (via interaction pathways, 𝑏_𝑖𝑗_, 𝑏_𝑖𝑘_); and (2) how coherent the environmental responses are across the community. Positive co-responses (𝑐𝑜𝑣(Δ𝑟_𝑘_, Δ𝑟_𝑙_) > 0) will amplify variance when interaction effects have the same sign, reinforcing each other. Conversely, negative co-responses buffer variance when interaction effects are aligned, as opposing environmental responses counteract their combined influence on species 𝑖.

When environmental co-responses are weak, they are unlikely to align with strong interaction effects, limiting the propagation of environmental variance through the network. But when co-responses are strong, even random alignment with key interaction pathways can amplify variance and destabilize community dynamics. Thus, the shape of the ENC distribution— whether clustered or skewed—can act as a proxy for community vulnerability, especially when interpreted alongside network structure to assess whether central species amplify or buffer environmental change.

### Concluding remarks and future perspectives

Ecological Network Coherence (ENC) formalizes the idea that variation in species’ environmental co-responses, while meaningful on their own, gains additional significance for anticipating community dynamics when viewed through interaction networks, which shape how responses propagate through the community. By capturing the distribution of co-responses among interacting species, ENC provides a general and flexible framework for quantifying how coherence is organized in ecological communities. We propose to focus on the types of data that are already broadly available for many taxa and ecosystems— occurrence records, abundance time series, and binary interaction networks—so that ENC can be readily applied and explored across species-rich systems. Our empirical applications illustrate its versatility: in a rainforest pollination network, ENC showed that interacting species tended to track temperature more coherently than the community at large, with highly connected pollinators emerging as potential stabilizers of coherence; in a collapsing marine fish community, ENC revealed how coherence was lost, fragmented, and re-concentrated through collapse and recovery. Although ENC distributions are defined as descriptive summaries of variability in co-responses, our Lotka–Volterra analyses and simulations demonstrate that their shapes can influence stability, suggesting that they hold promise as network-level indicators for anticipating disruptive ecological change.

Moving forward, two complementary research directions open to better understand the emergence and consequences of ENC. On the empirical side, central questions include: how coherent are ecological networks? and how does coherence vary with biogeography, community characteristics, and human impacts? One way to begin answering these questions is by describing ENC across multiple communities. This is now feasible thanks to available regional and global network databases of pollination (Lanuza et al. 2025), seed-dispersal (Mendes et al. 2024), terrestrial vertebrate trophic networks (Maiorano et al. 2010), and fish food webs (Albouy et al. 2019), that can be combined with species distribution models (SDMs) to estimate ENC in intrinsic responses to temperature. This would make it possible to identify hotspots of positive or negative coherence, to assess how coherence varies across latitude or with human footprint, and to examine whether network roles differ in their coherence—for example, whether super-generalist pollinators or predators remain aligned with their partners compared to specialists. A promising next step is to extend this approach into the future: by combining projected species distribution shifts and expected network rewiring under climate change (e.g. Albouy et al. 2014; Hao et al. 2025), it becomes possible to forecast how ENC might reorganize under global change.

Beyond intrinsic responses, the growing availability of abundance time series across taxa (Dornelas et al. 2018, Comte et al. 2021) enables estimation of ENC from population dynamics across multiple communities. Hierarchical models, such as the dynamic Bayesian framework outlined in Box 2, can then be used to recover latent variables and explore their link to potential drivers of repsonses such as climate or human pressures, providing a way to investigate what shapes ENC.

On the theoretical side, key questions are: what mechanisms drive the emergence of specific ENC distributions? and how can these distributions be used to anticipate disruptive community dynamics? Allometric trophic network models (Gauzens et al. 2023, Martinez 2023) provide a powerful framework for addressing these questions. They link body size to food-web structure, energy fluxes, and interaction strengths, while incorporating temperature dependence through Arrhenius functions (Brown et al. 2012). By tuning temperature-sensitive parameters to reflect different ENC distributions—that is, coherence in thermal responses among interacting species—we can ask how distributional shapes (clustered, skewed, polarized) influence stability and reorganization under warming (Petchey et al. 2010).

These predictions can be confronted with data: fish food webs are a particularly promising case, as they are strongly size-structured and highly temperature-sensitive (Brose et al. 2012). In addition, global datasets of traits, phylogeny, occurrences, and abundance make it possible to estimate their coherence to temperature at large scales (Albouy et al. 2019; Fernandes et al. 2020; Comte et al. 2021). Experimental systems offer a complementary approach by enabling direct manipulation of coherence and observation of its consequences for dynamics. In phytoplankton microcosms, already used to study abundance synchrony (Thompson et al. 2015), coherence structures could be altered by controlled perturbations such as warming. Tri-trophic insect–plant–parasite assemblages (Barbour et al. 2016; Barbour 2022; Gawecka et al. 2025) provide another tractable system, where different network architectures can be assembled to examine how structure shapes the effects of coherence on community dynamics. For instance, it could be interesting to explore whether networks with higher connectance are more vulnerable to propagating the potentially destabilizing effects of strong coherences (Martins et al. 2024).

Several challenges remain. First, computing the net effects of direct and indirect interactions across diverse taxa, trophic levels, and ecosystems is currently unfeasible with available data (Paine et al. 2018). An important next step is therefore to clarify how direct binary interactions, which define ENC, can be meaningfully connected to the net effects captured in Lotka–Volterra models. Second, species’ biological characteristics, such as their generation times, may further modulate coherence (Pureswaran et al. 2018, 2019). Some intuitive adjustments could help standardize comparisons across taxa to reflect these differences, such as weighting responses by body size, with smaller organisms expected to show stronger environmental sensitivities (Martin & Palumbi 1993). In addition, the current formulation assumes fixed networks and static responses, overlooking interaction rewiring (Bartley et al. 2019, Barbour & Gibert 2021) as well as evolutionary or plastic shifts in sensitivity (Urban et al. 2024). Similarly, fixed species pools omit turnover and colonization, which should be incorporated to extend ENC into more dynamic metacommunity contexts (Wang et al. 2019).

Despite these challenges, ENC provides a promising foundation for network-level biodiversity indicators. With continued development, it may help map ecosystems at risk, track community reorganization under climate change, and contribute to early-warning systems that complement existing indicators in global biodiversity frameworks.

## Acknowledgements

We thank Kim Gauthier and the Computational Biodiversity Science and Services (BIOS²) training program for supporting the organization of the working group that helped with developing this project, and the Université de Montréal for assistance with logistics. Financial support was provided by the NSERC - CREATE Training program in computational biodiversity science and the NSERC Discovery Grant to DG.

## Notes

### Competing Interest Statement

The authors have declared no competing interest.

