## Supplement 1 for "Tracking community change via network coherence"

### S1.1 Box 1

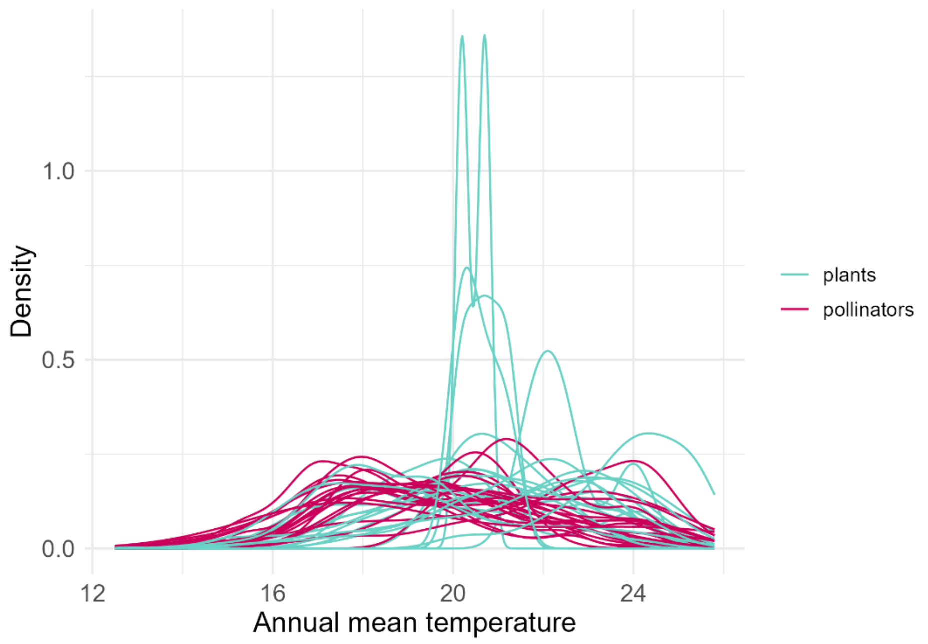


**Figure S 1. Species' suitability curves to temperature.**

Probability density functions for each species across the Atlantic Rainforest biome in Brazil, estimated from occurrence data.

### S1.2 Box 2

The dataset used in this box consists of the average yearly biomass for all trawls of 29 groundfish species in the North Atlantic Fishing Organization (NAFO) divisions 2J, 3K, and 3L, as published by Pedersen et al. (2017). Biomass was measured as kilograms per trawl of each species, which were recorded as part of a random depth-stratified survey with fixed duration and speed trawls (Pedersen et al. 2017). The dataset excludes species that were sensitive to a gear change from 1995 onwards, when the introduction of a smaller mesh size captured more small-bodied fish (Pedersen et al. 2017). To fit the model, we centered all species' biomasses on their biomass in 1981 and divided this difference in biomass by each species' mean biomass to model the normalized difference in each species' population size from the start of the time series.

We modelled fish biomass through time using a hierarchical model that shares information across populations to estimate their joint temporal trends while accounting for their covariation. We implemented this model as a Bayesian dynamic generalized additive model (DGAM) using the R package *mvgam*, which jointly estimates trends in multiple time series with a dynamic latent component that captures temporal associations between species (Clark & Wells 2023). The correlations between population trends are captured as species' relationships with latent temporal predictors, which allow information to be shared across populations at each time step to estimate the average trend. See Hébert (2024) for more details.

We specified the model as follows for species (j) and for time steps (t):

$$E\left(N_{i,t}\right) = \beta_{0,i} + \sum_{j=1}^{J} \beta_j s_j(t) + \sum_{m=1}^{M} \left(z_m, \lambda_{i,t}\right) + \varepsilon_{i,t} \quad \text{Equation S1}$$

where is the estimated biomass *N* for species *i* at time *t*, and is a species-specific intercept. The temporal component of the model consists of *j* smoothed basis functions at time *t*, which estimates the global temporal trend jointly for all species. The latent component of the model consists of $(z_m, \lambda_{i,t})$, which are estimates for M temporal latent predictors $(z_m)$, and $\lambda_{i,t}$, which are the species' latent factor loadings at each time $t$. To capture the latent temporal dynamics in populations' sizes, we specified the latent trend model as a smooth function in the form of a Gaussian process. The model estimated between M = 2 and M = 15 (half the total number of time series) latent predictors and increasingly penalized them to squeeze unneeded variables into flat lines, effectively removing them from the model (Clark & Wells, 2022). As a result of this procedure, the model retained M = 2 latent predictors.

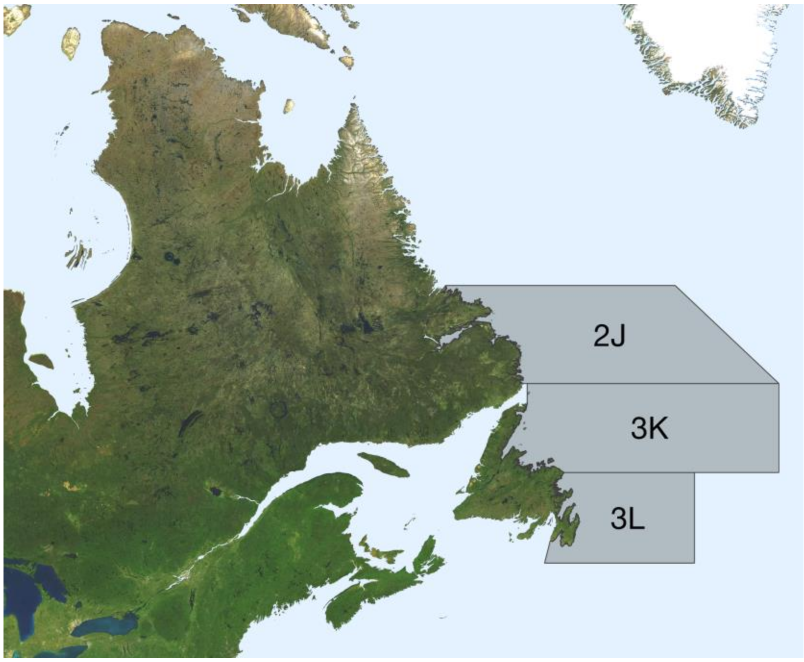


**Figure S 2. Study area for case study 2.**

The NAFO divisions (2J, 3K, 3L) of the Northwest Atlantic Ocean, where the groundfish community was monitored between 1981 and 2013.
